# Sleep duration and efficiency moderate the effects of prenatal and childhood ambient pollutant exposure on global white matter microstructural integrity in adolescence

**DOI:** 10.1101/2025.02.13.638133

**Authors:** Devyn L. Cotter, Orsolya Kiss, Hedyeh Ahmadi, Alethea de Jesus, Joel Schwartz, Fiona C. Baker, Daniel A. Hackman, Megan M. Herting

## Abstract

**Background:** Air pollution is a ubiquitous neurotoxicant associated with alterations in structural connectivity. Good habitual sleep may be an important protective lifestyle factor due to its involvement in the brain waste clearance and its bidirectional relationship with immune function. Wearable multisensory devices may provide more objective measures of sleep quantity and quality. We investigated whether sleep duration and efficiency moderated the relationship between prenatal and childhood pollutant exposure and whole-brain white matter microstructural integrity at ages 10-13 years.

**Methods:** We used multi-shell diffusion-weighted imaging data collected on 3T MRI scanners and objective sleep data collected with Fitbit Charge 2 from the 2-year follow-up visit for 2178 subjects in the Adolescent Brain Cognitive Development Study®. White matter tracts were identified using a probabilistic atlas. Restriction spectrum imaging was performed to extract restricted normalized isotropic (RNI) and directional (RND) signal fraction parameters for all white matter tracts, then averaged to calculate global measures. Sleep duration was calculated by summing the time spent in each sleep stage; sleep efficiency was calculated by dividing sleep duration by time spent in bed. Using an ensemble-based modeling approach, air pollution concentrations of PM_2.5_, NO_2_, and O_3_ were assigned to each child’s residential addresses during the prenatal period (9-month average before birthdate) as well as at ages 9- 10 years. Multi-pollutant linear mixed effects models assessed the associations between global RNI and RND and sleep-by-pollutant interactions, adjusting for appropriate covariates.

**Results:** Sleep duration interacted with childhood NO_2_ exposure and sleep efficiency interacted with prenatal O_3_ exposure to affect RND at ages 10-13 years. Longer sleep duration and higher sleep efficiency in the context of higher pollutant exposure was associated with lower RND compared to those with similar pollutant exposure but shorter sleep duration and lower sleep efficiency.

**Conclusions:** Low-level air pollution poses a risk to brain health in youth, and healthy sleep duration and efficiency may increase resilience to its harmful effects on white matter microstructural integrity. Future studies should evaluate the generalizability of these results in more diverse cohorts as well as utilize longitudinal data to understand how sleep may impact brain health trajectories in the context of pollution over time.

## Introduction

Ambient air pollutants are ubiquitous toxicants that pose a known risk to human health, and they have increasingly been linked to alterations in brain and mental health outcomes across the lifespan (1–4). The World Health Organization (WHO) and the United States Environmental Protection Agency (U.S. EPA) track numerous criteria pollutants, among them particulate matter with diameter <2.5 μm (PM_2.5_), nitrogen dioxide (NO_2_), and ground-level ozone (O_3_) (5). PM_2.5_ and NO_2_ are products of combustion of gasoline, oil, diesel fuel, coal, or wood, while ground-level O_3_ is produced via photooxidation of volatile organic compounds and other precursors by ultraviolet sunlight (6–8). When inhaled, all three pollutants may interact with the lung alveoli to induce an innate immune response, resulting in systemic circulation of cytokines, increased oxidative stress, and the weakening of tissue barriers such as the nasal epithelium, blood-brain barrier (BBB), and the blood-placental barrier (9–11). It is thought that children are particularly susceptible to air pollution-related harm because they have higher respiratory rates, higher rates of neurodevelopmental change, and increased time spent outside compared to adults (12,13). Timing of exposure (i.e., prenatal versus childhood) as well as individual factors like sex may contribute to differential mechanisms by which air pollution increases risk for various diseases or disorders (1,14,15).

The brain connectome is defined as the spatial map of neural connections that underlie all motor, cognitive, emotional, and behavioral functions (16). Structural connectivity is characterized by white matter microstructural integrity of tracts connecting various brain regions. Air pollution exposure during development has increasingly been associated with changes in structural connectivity, both cross-sectionally and over time (2). Using data from the nationwide Adolescent Brain and Cognitive Development (ABCD) Study in the United States, our group has led multiple studies investigating the link between pollutant exposure and white matter microstructural integrity as measured using restriction spectrum imaging (RSI), an advanced multi-compartment diffusion model that can differentiate between extracellular and intracellular directional and isotropic diffusion (17–19). The first cross-sectional analysis found a positive association between childhood PM_2.5_ exposure and intracellular, restricted isotropic diffusion (RNI) at ages 9-10 years old, suggestive of a change in glial cell morphology or quantity which we hypothesized may reflect neuroinflammation. Next, we conducted a longitudinal study that included childhood exposure to three criteria pollutants (i.e., annual average daily PM_2.5_, daily NO_2_, daily 8-hour maximum O_3_) and found that higher childhood NO_2_ exposure at ages 9-10 years was associated with attenuated longitudinal increases of RNI throughout the brain in female youth from ages 9-13 years-old (19). In contrast, we found higher childhood O_3_ exposure had similar effects on RNI in both sexes from ages 9-13 years, albeit more strongly in males (19). In a follow-up sex-stratified multivariate cross-sectional analysis at ages 10-13 years, we expanded this research to also include prenatal exposure to PM_2.5_, NO_2_, and O_3_, alongside childhood exposure on white matter microstructure (18). We found prenatal and childhood exposure positively correlated with RNI as well as intracellular, restricted directional (RND) diffusion in white matter in female youth, but negatively correlated with the same metrics in male youth, with the impacted tracts varying by sex (18). Additionally, using diffusion tensor imaging (DTI) data from the Generation R study, a large Netherlands-based birth cohort, researchers found that both prenatal and childhood (0-4 years-old) exposure to PM_2.5_ and its components, NO_2_, and nitrogen oxides (NO_X_) were linked to lower fractional anisotropy (FA) and higher mean diffusivity (MD) throughout the brain at ages 9-12 years (20,21). Recent work in the Generation R cohort examined the longitudinal associations between prenatal and childhood exposure to multiple pollutants and white matter DTI measures in children aged 9-17 years (median age 9.9 years) over two time points (22). They found that prenatal exposure to PM_2.5_ and childhood exposure to PM (size fractions 10, 2.5, 2.5-10 ug/m^3^) and NO_X_ was related to lower global FA. Additionally, prenatal exposure to silicon (a component of PM_2.5_) and the oxidative potential of PM_2.5_ as well as childhood exposure of PM_2.5_ was associated with accelerated decreases of MD over time. In another DTI study, Peterson and colleagues (23) found that exposure to higher PM_2.5_ during gestation was linked to a higher average diffusion coefficient in large posterior white matter fiber bundles – indicative of reduced myelin and/or fiber density/coherence. However, pollutant exposure was not associated with white matter FA in youth aged 6-14 years. This suggests that increased pollutant exposure during various windows of pre- and postnatal development are cross-sectionally associated with reduced white matter microstructural integrity in late childhood to early adolescence, but both accelerated (i.e., faster MD decreases (22) and faster RND increases (19)) and attenuated (i.e., slower RNI increases (19)) white matter microstructural development over time, depending on the diffusion metric utilized. Considering this compelling evidence that air pollution during vulnerable pre- and postnatal windows of development may alter brain connectivity, as well as studies that suggest air pollution is linked to poor mental health outcomes and neurodevelopmental disorders (24), it is important to understand if individual differences in lifestyle factors may contribute to resilience in the face of harmful environmental exposures.

Potential protective factors that may moderate air pollution’s negative effects on brain outcomes include quantity and quality of sleep. Sleep is well-known to be highly correlated with the immune system in a bidirectional manner to maintain the body’s homeostasis and support cognitive and emotional functions important for everyday life (25). When one system is dysregulated, the negative effects can reverberate, affecting multiple biological systems and outcomes including the brain. Animal studies have found that cytokines and prostaglandins play a crucial role in regulating sleep-wake cycles (25). In fact, disruptions in prostaglandin levels have been associated with sleep disturbances such as decreased efficiency and increased overnight awakenings, as well as decreased slow-wave sleep (25). Though the exact mechanisms are not well understood, sufficient sleep has been shown to restore normal levels of upregulated immune cell populations and improve adaptive immune responses (25). While much remains to be discovered in sleep-immune crosstalk, the current literature robustly supports the notion that sleep is integral in proper immune function and overall health and wellbeing. As air pollution is known to induce aberrant systemic immune activity with potential to induce neuroinflammation (1,26), sleep’s role in immune function may provide a pathway for sleep quantity and quality to protect the brain against the neurotoxic effects of air pollution exposure. To this end, in the first study of its kind, sleep quality was recently shown to mitigate the negative effects of air pollution on biological aging in a stepwise manner in an adult human sample from the UK Biobank, such that accelerations in biological aging associated with air pollution exposure were significantly slowed by higher sleep efficiency (27). Yet, similar questions have not yet been explored in adolescent populations or pertaining to brain health specifically.

Leveraging data from 2178 subjects enrolled in the ABCD Study, the current cross-sectional study aimed to examine the potential moderating effect of sleep duration and efficiency measured with a wrist-worn commercial device (Fitbit Charge 2) on the relationship between pollutant exposure during two developmental windows (i.e., prenatal and childhood) and white matter microstructural integrity in youths aged 10-13 years. Additionally, due to sex-specific effects in environmental neurotoxicity (28), brain development (29), and measures of sleep health (30), we also investigated potential sex differences in how sleep may mitigate the negative effects of air pollution on structural brain connectivity. Because of potential opposing effects of air pollution and sleep on biological functions, such as immune health, we hypothesized that longer sleep duration as well as better sleep efficiency would diminish the negative effects of air pollution exposure on global white matter microstructural integrity in adolescence. The results discussed here suggest that sleep may protect young brains against the neurotoxic effects of air pollution.

## Methods

### Study Population

The ABCD Study® is a large and regionally diverse study of neurodevelopment in youth from 21 communities across the United States. Between the years 2016 to 2018, 11,876 children between the ages of 9-10 years were enrolled, with plans to follow them annually over the course of 10 years into young adulthood (31). An overview of detailed recruitment procedures have been previously described (32). The ABCD Study’s inclusion criteria included age (9-10 years old at initial visit) and English language proficiency. Exclusion criteria were as follows: major medical or neurological conditions, history of traumatic brain injury, diagnosis of schizophrenia, moderate/severe autism spectrum disorder, intellectual disability, alcohol/substance use disorder, premature birth (gestational age <28 weeks), low birthweight (<1200 g), and contraindications to magnetic resonance imaging (MRI) scanning. The ABCD Study obtained approval for all study procedures from the University of California, San Diego centralized institutional review board (IRB# 160091). Subsequently, each study site was also required to obtain approval from their respective institutional review boards. All parents or caregivers provided written informed consent and children provided written assent.

Data used in the current analyses were obtained from the ABCD’s 5.0 Data Release (33). 2178 subjects from 21 sites across the U.S. were included (Supplemental Figure 2). Due to the availability of wrist wearable data from the Fitbit Charge 2 at the 2-year follow-up visit only, we used cross-sectional wrist wearable and neuroimaging data from the 2-year follow-up visit when subjects were aged 10-13 years. All subjects had air pollution concentration estimates from the prenatal and childhood (ages 9-10 years, baseline visit) periods, as well as high quality MRI scans without incidental findings of clinical significance and wrist wearable data collected within the protocol period (see below for quality control details). MRI scans were collected on Siemens Prisma, Philips, or GE 750 3T MRI scanners using harmonized acquisition procedures specific to the ABCD Study, as previously described by Casey et al. (34). Importantly, the final sample used here excluded participants with neuroimaging and wrist wearable data collected after March 1, 2020, so as to remove any potential confounding effects of the COVID-19 pandemic, an event that significantly disrupted normal routines and increased perceived stress (35). Please see Table 1 for detailed cohort characteristics.

**Table 1.** Cohort demographic and socioeconomic characteristics, pollutant levels, and sleep metrics.

| Cohort Characteristics |  |
| --- | --- |
| Total N | 2178 |
| Sex [F], N (%) | 995 (45.7%) |
| Mean Age [months], (SD) | 143.12 (7.72) |
| Pubertal Development Scale, N (%) |  |
| 1 (pre-pubertal) | 534 (24.5%) |
| 2 (early puberty) | 550 (25.3%) |
| 3 (mid-puberty) | 734 (33.7%) |
| 4 (late puberty) | 340 (15.6%) |
| 5 (adult-like) | 20 (0.9%) |
| Race/Ethnicity, N (%) |  |
| Non-Hispanic White | 1424 (65.4%) |
| Non-Hispanic Black | 144 (6.6%) |
| Hispanic | 367 (16.8%) |
| Non-Hispanic Asian* | 38 (1.7%) |
| Multi-Racial/Other*† | 205 (9.4%) |
| Highest Household Education, N (%) |  |
| Post Graduate Degree | 887 (40.7%) |
| Bachelor | 660 (30.3%) |
| Some College | 480 (22.1%) |
| HS Diploma/GED | 111 (5.1%) |
| < HS Diploma | 40 (1.8%) |
| Overall Income (USD), N (%) |  |
| ≥100K | 1057 (48.5%) |
| ≥50K & <100K | 624 (28.7%) |
| <50K | 384 (17.6%) |
| Don't Know/Refuse | 113 (5.2%) |
| Mean Pollutant Levels (SD) |  |
| Prenatal PM<sub>2.5</sub>, $\mu\text{g}/\text{m}^3$ | 10.78 (2.42) |
| Childhood PM<sub>2.5</sub>, $\mu\text{g}/\text{m}^3$ | 7.31 (1.58) |
| Prenatal NO<sub>2</sub>, ppb | 25.59 (10.18) |
| Childhood NO<sub>2</sub>, ppb | 18.01 (5.98) |
| Prenatal O<sub>3</sub>, ppb | 40.06 (4.69) |
| Childhood O<sub>3</sub>, ppb | 41.37 (4.29) |
| Mean Fitbit Charge 2 Sleep Variables (SD) |  |
| Sleep Duration (hours) | 7.51 (0.64) |
| Sleep Efficiency (%) | 0.87 (0.02) |

### Ambient Air Pollution Estimates

Geocoded information about participants’ residential addresses was used to define the locations where prenatal and one-year childhood exposures to PM_2.5_, NO_2_, and O_3_ were estimated (36). Primary residential addresses at study enrollment (i.e., when the child was 9-10 years) were collected in-person from the participant’s caregiver during the study visit between October 2016 to October 2018. At the 2-year follow-up visit, additional previous residential addresses were collected retrospectively via caregiver report. All residential addresses were geocoded by the ABCD consortium’s Data Analytics Information and Resource Center (DAIRC) (36). Daily ambient air pollution concentration estimates for PM_2.5_, NO_2_, and 8-hour maximum O_3_ were then estimated for the entire continental U.S. as previously described (36). Briefly, hybrid spatiotemporal models were leveraged to first derive daily air pollution estimates at a 1-km^2^ resolution, utilizing satellite remote sensing, land-use regression, and chemical transport models (36–38). Daily estimates were subsequently averaged over the 2016 calendar year, corresponding with participant study enrollment when children were aged 9-10 years. One-year annual average concentrations during childhood were then assigned to primary residential addresses for each participant. To estimate prenatal exposure, daily exposure estimates for 9 months of pregnancy based on the child’s birthdate [birth years 2005-2009] we averaged and assigned to the address that corresponded to the child’s birth year. If multiple addresses overlapped with the child’s birthdate, the prenatal average exposure values for each residence were weighted by the reported percent of time spent at that residence, after which the sum of these weighted exposure averages was divided by the sum of all reported percentages. To reduce potential misclassification bias, subjects were excluded from the analyses if the percentage of time reported across the multiple addresses overlapping with the child’s birthdate totaled below 90% or above 110%. Quality-controlled prospective residential addresses (i.e., at time 1- or 2-year follow-up) are not currently available within the ABCD dataset. Thus, we assumed the spatial contrast remained constant between the study enrollment period and the annual 2-year follow-up visit, as demonstrated using these ensemble-based models from 2000 to 2016 (37–39). In our final models, we also covaried for those that had moved locations since the baseline visit. Lastly, standardized pollutant values were obtained by subtracting the mean and dividing by 5 for each pollutant.

### Wearable Technology Measures of Sleep

Given that subjective measures of sleep quantity and quality can be biased by self-reporter error, objective measurement of sleep with a wearable device represents a non-invasive way to estimate sleep parameters more accurately. Polysomnography, including electroencephalogram (EEG), electro-oculogram, electromyogram, electrocardiogram, pulse oximetry, and airflow/respiratory effort, remains the gold standard in sleep research for objectively measured sleep, but a recent study indicated that there was substantial agreement between Fitbit and home-based EEG methods in measuring total sleep duration (26). Thus, we examined objective measures of sleep, collected from a Fitbit Charge 2 device. Adolescents wore the device for three consecutive weeks starting after their annual visit at the 2-year follow-up (40). A valid week was defined as at least 4 days of sleep data including at least one weekend day (40). Subjects were included if they had at least one valid week collected within the protocol period. Parameters of interest included total sleep duration (*hours*) and sleep efficiency (*percent*). Total sleep duration was calculated by summing time spent in light, deep, and REM stages, to account for overnight awakenings. Sleep efficiency was calculated by dividing sleep duration by time in bed. Time in bed was defined as the difference between the time of day the participant got out of bed in the morning and the time of night they went to bed the night before, but were not necessarily asleep, as determined by Fitbit. Weekly weighted averages of sleep duration and efficiency were calculated and used in the final models.

### Restriction Spectrum Imaging (RSI)

Multi-shell diffusion-weighted images were acquired using multiband echo-planar imaging (41,42) with slice acceleration factor 3 and a 1.7 mm^3^ resolution, alongside a fieldmap scan for B0 distortion correction. Diffusion weights included seven b=0 frames and 96 total diffusion directions at 4 b-values, with 6 at bl=l500ls/mm^2^, 15 at bl=l1000ls/mm^2^, 15 at bl=l2000ls/mm^2^, and 60 at bl=l3000ls/mm^2^ (43). Following distortion, bias field, and motion correction, manual and automated quality control were conducted on all images (43). Using this multi-shell sequence, RSI allows for biophysical modeling of both intra- and extracellular compartments of tissue within the brain (44). Selected RSI model outputs are unitless on a scale of 0-1 and included both restricted (intracellular) normalized isotropic (RNI) and directional (RND) signal fractions of white matter fiber tract regions of interest (ROIs) created with AtlasTrack (45). RNI measures intracellular diffusion in all directions and likely represents diffusion within support cells or other round structures, while RND measures intracellular diffusion in a single direction and likely represents diffusion along an axon or other elongated process (44,46). Brain images were included if deemed absent of clinically significant incidental findings and passed all ABCD quality-control parameters. Given our previous whole brain findings between air pollution and structural connectivity (18,19), parameters of interest included global RND and global RNI, averaged across all AtlasTrack fibers.

### Confounders and Covariates

Time-invariant covariates were taken from enrollment at the baseline visit, and included race and ethnicity (*Asian, Hispanic, non-Hispanic Black, non-Hispanic White [reference group],* or *Multi-Racial/Other*), total household income in United States dollars (USD) (≥*100K, 100-50K, <50K [reference group],* or *Don’t Know/Refuse to Answer*), and highest household education (*Post-Graduate, Bachelor, Some College, High School Diploma/GED,* or *<High School Diploma [reference group]*). Race/ethnicity and socioeconomic factors were included because pollution levels are higher in minority communities and those from disadvantaged social status backgrounds (47). We also included the participant’s age (*months*), sex assigned at birth (*male, female*), and pubertal development stage (PDS; *1-5*, consistent with Tanner-like categorization

(48)) as subject-specific precision variables. MRI-specific precision variables included scanner manufacturer (*Siemens*, *Philips*, *GE [reference group]*) to account for differences in both scanner hardware and software, and average framewise displacement (*mm*) to account for head motion. Lastly, we covaried for season of visit (*Fall [reference group], Winter, Spring, Summer)*, given the seasonality in pollutant exposure concentrations, as well as whether participants moved in between the 2-year follow-up visit and the initial visit when childhood pollutant concentrations were measured. Supplemental Table 1 shows the comparison between the characteristics of the current study sample and the larger ABCD Study cohort.

### Statistical Analyses

We used hierarchical linear mixed-effect models, as implemented in *lme4*::*lmer()* (49) in R statistical software (Version 4.1.2.) (50) to account for the multi-level data structure, including random effects of family nested within study sites. Given our previous findings showing notable sex-specific effects in air pollution and brain outcomes (18,19), we examined sex differences in the moderating effect of total sleep duration (*hours*) on the relationship between exposure to pollutants (prenatal and childhood PM_2.5_, NO_2_, and O_3_) and brain outcomes (global RNI and RND) with a three-way pollutant-by-sleep-by-sex interaction term (which included three additional two-way interaction terms [pollutant-by-sleep, sex-by-sleep, pollutant-by-sex]). For model parsimony and ease of interpretation, the highest order interaction term (i.e., three-way pollutant-by-sleep-by-sex interaction term) was dropped if not significant at the level of *p*<0.05. Similar analyses were conducted for sleep efficiency (*percent*). For models demonstrating a significant relationship between the pollutant-by-sleep interaction term and global RNI or RND, we completed post-hoc analyses to determine if any specific tracts were primarily affected.

To account for co-exposure of the three criteria pollutants at two developmental windows, we controlled for the other pollutants not included in the interaction terms of interest, in addition to all covariates discussed above. Upon checking model assumptions, we found a violation of the heteroscedasticity assumption due to the inclusion of siblings from the same family. Therefore, we applied robust variance estimations (RVE) to all models to obtain reliable standard errors and test statistics, ensuring the robustness of our findings. This allowed for the preservation of the hierarchical data structure with fidelity to ABCD’s original study design. Given our hypotheses, we did not correct for multiple comparisons for the two outcomes of interest (i.e., global RNI and RND); however, a false discovery rate (FDR) adjustment was performed on post-hoc analyses examining each tract separately. For the models with significant pollutant-by-sleep interaction terms, we further probed the interaction by performing pairwise tests using the *emmeans::emmeans()* function in R (51).

## Results

We analyzed 2178 unique ABCD Study participants (45.7% female) from 21 sites throughout the U.S. to determine if sleep duration and efficiency moderated the relationship between prenatal and childhood exposure to three criteria pollutants (PM_2.5_, NO_2_, and O_3_) and white matter microstructural integrity in youths aged 10-13 years. Prenatal exposure estimates were higher than childhood exposure estimates for PM_2.5_ and NO_2_, but not for O_3_ (Table 1). Spearman correlations between pollutants from both developmental windows can be found in Supplemental Table 1. Overall, PM_2.5_ and O_3_ were negatively correlated (r_S_ ranges from -0.07 to -0.15), while PM_2.5_ and NO_2_ (r_S_ ranged from 0.16 to 0.31) as well as NO_2_ and O_3_ (r_S_ ranges from 0.04 to 0.15) were positively correlated (Supplemental Figure 1). Additionally, sleep duration and sleep efficiency were weakly positively correlated (r_S_ = 0.06) (Supplementary Figure 1). Of note, from a clinical standpoint, average sleep duration is low, with an average of 7.51 hours per night (*t*l=l -109.61 (*µ*l=l9), dfl=l 2177, *p*l=l0) (52). Average sleep efficiency is normal at 87% in our sample, with ≥85% sleep efficiency deemed acceptable across all age groups (53).

Across all models, the highest order interaction term (e.g., three-way pollutant-by-sleep-by-sex interaction term) did not demonstrate a significant relationship with any brain outcome (global RND and RNI) and thus was dropped for model parsimony and ease of interpretation. The lack of significance here indicates that there were no observed sex differences in how sleep metrics moderated the relationship between air pollution exposure and global white matter microstructural integrity. The following results are from simplified models.

### Moderating effect of total sleep duration on the association between air pollutants and structural brain connectivity at ages 10-13 years

Total sleep duration moderated the association between childhood NO_2_ exposure and global RND (b = -0.001, p = 0.006) (Table 2, Figure 1). Post-hoc pairwise tests demonstrated that there were no statistically significant associations between childhood NO_2_ and RND at 6, 7, or 8 hours of sleep duration; however, pairwise contrasts showed that sleep duration and childhood NO_2_ exposure significantly interacted to affect global RND, such that a cross-over effect was observed (Figure 1) and the slopes per level of sleep duration were significantly different from each other (p = 0.03), but not from zero (Supplemental Table 2). Post-hoc regional analyses of each separate tract revealed this association was strongest for the corpus callosum (b = -0.002, p_FDR_ = 0.0006) and right uncinate fasciculus (b = -0.001, p_FDR_= 0.003) (Supplemental Table 4).

**Figure 1.**
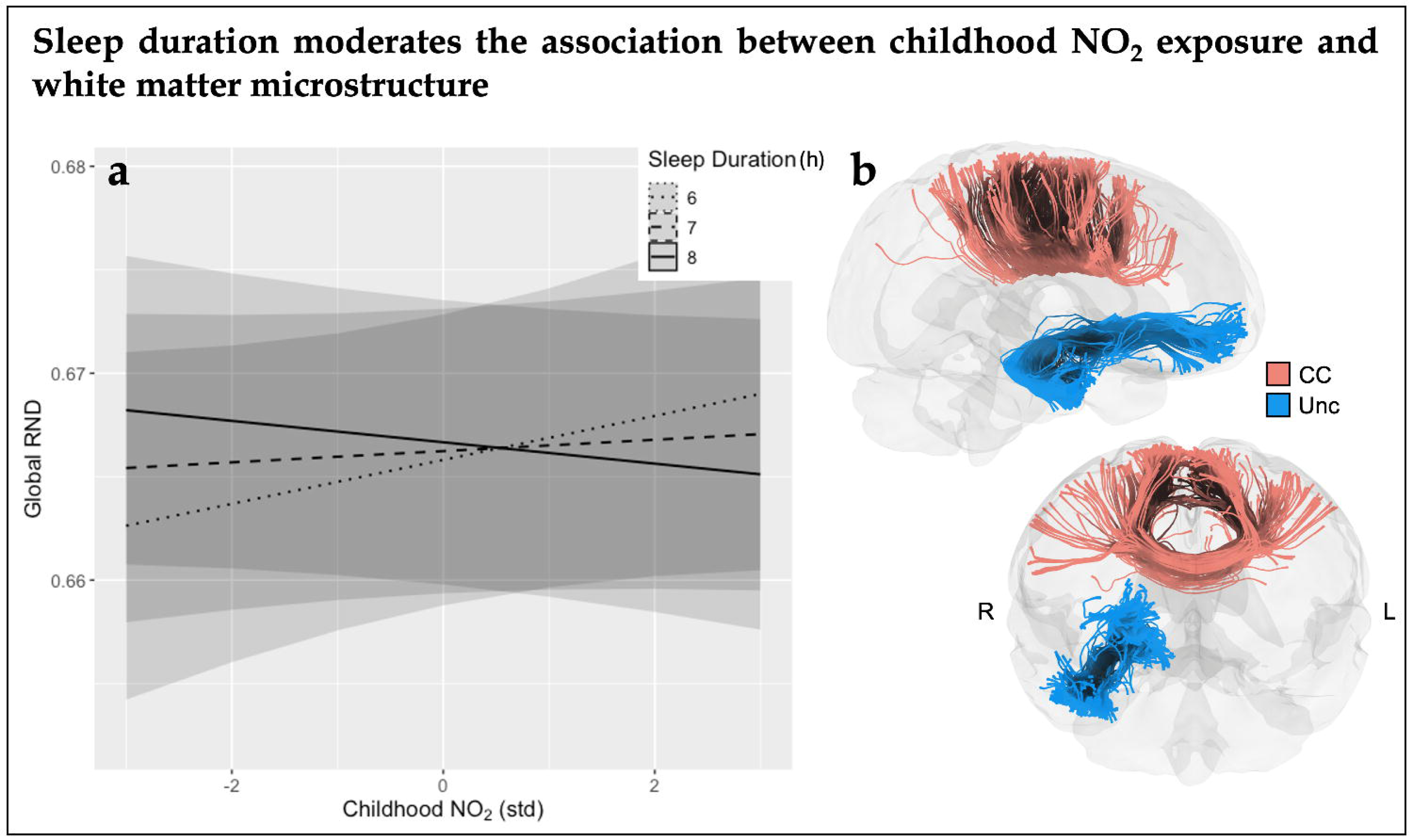
A) Significant interaction between childhood NO_2_ exposure and sleep duration on global intracellular directional diffusion (RND). Childhood NO_2_ is standardized, with 0 equal to the mean in our sample (18.01 *ppb*), and 1 unit representing a 5-*ppb* change. Sleep duration is presented in *hours*. B) Visualization of the individual tracts affected by the pollutant-by-sleep interaction term in the post-hoc regional analyses. Abbreviations: parts per billion (ppb), intracellular directional diffusion (RND), standardized (std), corpus callosum (CC), uncinate fasciculus (Unc), right (R), left (L).

**Table 2.** Results from multi-pollutant models examining how sleep duration interacts with pollutants to affect brain connectivity, including unstandardized betas, standard error (SE), 95% confidence intervals (CI), and p-values. Significant models are **bolded** (p < 0.05). Models were adjusted for pollutants not included in the interaction term, demographic and socioeconomic variables for each child, and precision MRI variables (see Methods). Abbreviations: intracellular isotropic diffusion (RNI), intracellular directional diffusion (RND), standardized (std), standard error (SE), confidence interval (CI).

| *Parameter* |  | Global RNI |  |  |  | Global RND |  |  |  |
| --- | --- | --- | --- | --- | --- | --- | --- | --- | --- |
|  |  | *Coefficient* | *SE* | *95% CI* | *p* | *Coefficient* | *SE* | *95% CI* | *p* |
| Prenatal | PM<sub>2.5</sub> (std) | 0.0056 | 0.0047 | -0.0036, 0.0149 | 0.231 | **0.0155** | **0.0073** | 0.0012, 0.0297 | **0.033** |
|  | Sleep Duration | -0.0002 | 0.0004 | -0.0009, 0.0006 | 0.649 | 0.0005 | 0.0005 | -0.0005, 0.0015 | 0.373 |
|  | PM<sub>2.5</sub> (std) x Sleep Duration | -0.0007 | 0.0006 | -0.0019, 0.0006 | 0.295 | -0.0018 | 0.001 | -0.0037, 0 | 0.053 |
|  | NO<sub>2</sub> (std) | -0.0011 | 0.0013 | -0.0036, 0.0014 | 0.398 | 0.0013 | 0.001 | -0.0007, 0.0034 | 0.205 |
|  | Sleep Duration | -0.0002 | 0.0004 | -0.001, 0.0006 | 0.636 | 0.0004 | 0.0005 | -0.0005, 0.0013 | 0.363 |
|  | NO<sub>2</sub> (std) x Sleep Duration | 0.0001 | 0.0002 | -0.0002, 0.0004 | 0.521 | -0.0002 | 0.0001 | -0.0005, 0.0001 | 0.166 |
|  | O<sub>3</sub> (std) | -0.0009 | 0.0032 | -0.0071, 0.0053 | 0.778 | -0.0018 | 0.0045 | -0.0107, 0.0071 | 0.698 |
|  | Sleep Duration | -0.0002 | 0.0004 | -0.001, 0.0006 | 0.646 | 0.0004 | 0.0005 | -0.0005, 0.0013 | 0.351 |
|  | O<sub>3</sub> (std) x Sleep Duration | 0.0002 | 0.0004 | -0.0006, 0.001 | 0.670 | 0.0004 | 0.0006 | -0.0008, 0.0015 | 0.550 |
| Childhood | PM<sub>2.5</sub> (std) | 0.0044 | 0.0096 | -0.0143, 0.0232 | 0.643 | 0.015 | 0.0132 | -0.011, 0.0409 | 0.258 |
|  | Sleep Duration | -0.0002 | 0.0004 | -0.001, 0.0006 | 0.641 | 0.0005 | 0.0005 | -0.0005, 0.0015 | 0.365 |
|  | PM<sub>2.5</sub> (std) x Sleep Duration | -0.0004 | 0.0013 | -0.0029, 0.0022 | 0.775 | -0.002 | 0.0017 | -0.0053, 0.0013 | 0.240 |
|  | NO<sub>2</sub> (std) | 0.0013 | 0.0013 | -0.0012, 0.0038 | 0.317 | **0.0058** | **0.0023** | 0.0013, 0.0103 | **0.011** |
|  | Sleep Duration | -0.0002 | 0.0004 | -0.001, 0.0006 | 0.631 | 0.0004 | 0.0005 | -0.0005, 0.0014 | 0.371 |
|  | NO<sub>2</sub> (std) x Sleep Duration | -0.0002 | 0.0002 | -0.0005, 0.0002 | 0.325 | **-0.0008** | **0.0003** | -0.0013, -0.0002 | **0.006** |
|  | O<sub>3</sub> (std) | -0.0017 | 0.0037 | -0.009, 0.0056 | 0.651 | -0.0045 | 0.0053 | -0.0149, 0.0059 | 0.399 |
|  | Sleep Duration | -0.0002 | 0.0004 | -0.001, 0.0006 | 0.648 | 0.0004 | 0.0005 | -0.0006, 0.0014 | 0.388 |
|  | O<sub>3</sub> (std) x Sleep Duration | 0.0002 | 0.0005 | -0.0008, 0.0012 | 0.753 | 0.0005 | 0.0007 | -0.0009, 0.0019 | 0.503 |

There were no other statistically significant interactions between other air pollutant exposures and sleep duration on global RND or RNI. There was a significant main effect between prenatal PM_2.5_ exposure and global RND (b = 0.02, p = 0.03), but no other significant main effects of pollutants or sleep duration on global RNI or RND. All results can be found in Table 2.

### Moderating effect of sleep efficiency on the association between air pollutants and structural brain connectivity at ages 10-13 years

Sleep efficiency moderated the association between prenatal O_3_ and global RND (b = -0.03, p = 0.03) (Table 3, Figure 2). Post-hoc pairwise tests demonstrated that the relationship between prenatal O_3_ exposure and global RND was positive and statistically significant at the first quantile (86%) and median sleep efficiency levels (87%), with the slope diminishing as sleep efficiency rose; at the third quantile of sleep efficiency (88%), there was no relationship between prenatal O_3_ exposure and global RND (Supplemental Table 3). All pairwise contrasts showed statistically significant differences in trends at different levels of sleep efficiency, with stronger trends at lower levels of sleep efficiency (86%, 87%) (Supplemental Table 3). This indicates that higher sleep efficiency reduced the association between prenatal O_3_ exposure and RND. Post-hoc regional analyses of each separate tract revealed this association was strongest for the right corticospinal tract (b = -0.04, p_FDR_ = 0.009) (Supplemental Table 5).

**Figure 2.**
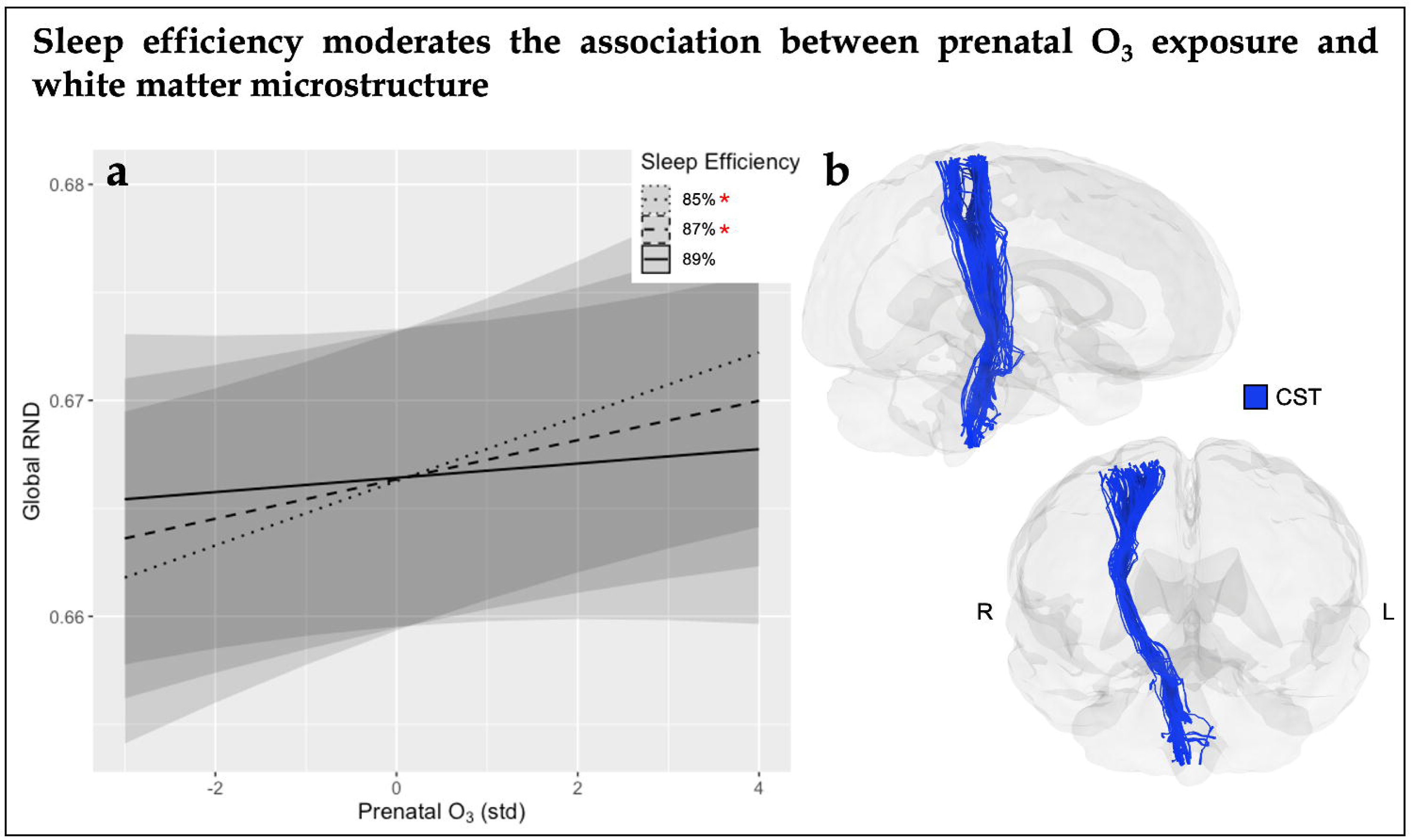
A) Significant interaction between prenatal O_3_ exposure and sleep efficiency on global intracellular directional diffusion (RND). Prenatal O_3_ is standardized, with 0 equal to the mean in our sample (40.06 *ppb*), and 1 unit representing a 5-*ppb* change. Sleep efficiency is presented in *percentage*. Red asterisks represent statistically significant slopes. B) Visualization of the individual tract affected by the pollutant-by-sleep interaction term in the post-hoc regional analyses. Abbreviations: parts per billion (ppb), intracellular directional diffusion (RND), standardized (std), corticospinal tract (CST), right (R), left (L).

**Table 3.** Results from multi-pollutant models examining how sleep efficiency interacts with pollutants to affect brain connectivity, including unstandardized betas, standard error (SE), 95% confidence intervals (CI), and p-values. Significant models are **bolded** (p < 0.05). Models were adjusted for pollutants not included in the interaction term, demographic and socioeconomic variables for each child, and precision MRI variables (see Methods). Abbreviations: intracellular isotropic diffusion (RNI), intracellular directional diffusion (RND), standardized (std), standard error (SE), confidence interval (CI).

| *Parameter* |  | Global RNI |  |  |  | Global RND |  |  |  |
| --- | --- | --- | --- | --- | --- | --- | --- | --- | --- |
|  |  | *Coefficient* | *SE* | *95% CI* | *p* | *Coefficient* | *SE* | *95% CI* | *p* |
| Prenatal | PM<sub>2.5</sub> (std) | 0.0107 | 0.0197 | -0.0279, 0.0494 | 0.586 | 0.0114 | 0.0258 | -0.0393, 0.062 | 0.659 |
|  | Sleep Efficiency | -0.005 | 0.012 | -0.0286, 0.0186 | 0.679 | 0.0044 | 0.013 | -0.021, 0.0299 | 0.733 |
|  | PM<sub>2.5</sub> (std) x Sleep Efficiency | -0.0116 | 0.0231 | -0.0569, 0.0338 | 0.617 | -0.0114 | 0.03 | -0.0703, 0.0475 | 0.705 |
|  | NO<sub>2</sub> (std) | 0.0056 | 0.0055 | -0.0051, 0.0163 | 0.307 | 0.0046 | 0.0058 | -0.0067, 0.0159 | 0.425 |
|  | Sleep Efficiency | -0.0066 | 0.0126 | -0.0312, 0.0181 | 0.602 | 0.0031 | 0.0135 | -0.0234, 0.0297 | 0.817 |
|  | NO<sub>2</sub> (std) x Sleep Efficiency | -0.0068 | 0.0062 | -0.0189, 0.0054 | 0.274 | -0.0054 | 0.0064 | -0.0179, 0.0071 | 0.398 |
|  | O<sub>3</sub> (std) | 0.0076 | 0.0156 | -0.0231, 0.0382 | 0.628 | **0.0261** | **0.0118** | 0.0031, 0.0492 | **0.026** |
|  | Sleep Efficiency | -0.0054 | 0.0123 | -0.0296, 0.0188 | 0.662 | 0.0038 | 0.011 | -0.0177, 0.0254 | 0.727 |
|  | O<sub>3</sub> (std) x Sleep Efficiency | -0.0082 | 0.018 | -0.0435, 0.027 | 0.647 | **-0.029** | **0.0135** | -0.0554, -0.0026 | **0.032** |
| Childhood | PM<sub>2.5</sub> (std) | 0.022 | 0.0278 | -0.0326, 0.0766 | 0.430 | 0.0494 | 0.0413 | -0.0317, 0.1305 | 0.232 |
|  | Sleep Efficiency | -0.0048 | 0.0122 | -0.0286, 0.019 | 0.692 | 0.0055 | 0.0119 | -0.0178, 0.0288 | 0.642 |
|  | PM<sub>2.5</sub> (std) x Sleep Efficiency | -0.0233 | 0.0321 | -0.0863, 0.0397 | 0.468 | -0.0567 | 0.0467 | -0.1484, 0.0349 | 0.225 |
|  | NO<sub>2</sub> (std) | 0.0051 | 0.0081 | -0.0109, 0.021 | 0.533 | 0.0137 | 0.0119 | -0.0098, 0.0371 | 0.253 |
|  | Sleep Efficiency | -0.006 | 0.0121 | -0.0298, 0.0177 | 0.618 | 0.0023 | 0.014 | -0.0253, 0.0298 | 0.873 |
|  | NO<sub>2</sub> (std) x Sleep Efficiency | -0.0058 | 0.0094 | -0.0242, 0.0126 | 0.537 | -0.0159 | 0.0138 | -0.0429, 0.0112 | 0.251 |
|  | O<sub>3</sub> (std) | -0.0003 | 0.0133 | -0.0264, 0.0258 | 0.985 | -0.0209 | 0.0144 | -0.0491, 0.0073 | 0.146 |
|  | Sleep Efficiency | -0.0054 | 0.0126 | -0.03, 0.0193 | 0.668 | 0.0061 | 0.0112 | -0.0158, 0.0281 | 0.585 |
|  | O<sub>3</sub> (std) x Sleep Efficiency | -0.0003 | 0.0152 | -0.0301, 0.0296 | 0.987 | 0.0231 | 0.0166 | -0.0094, 0.0556 | 0.163 |

There were no other statistically significant interaction effects seen between any other exposures and sleep efficiency on global RND. Lastly, there were no statistically significant main effects of pollutant or sleep efficiency on global RNI or RND. All results can be found in Table 3.

## Discussion

To our knowledge, this is the first study to investigate whether metrics of habitual sleep may moderate the association between air pollution exposure and white matter microstructure in adolescents. In testing the pollutant-by-sleep interaction terms, we found that sleep duration interacted with childhood NO_2_ exposure and sleep efficiency interacted with prenatal O_3_ exposure to affect global white matter intracellular directional diffusion at ages 10-13 years. We demonstrated that there were no significant effects of childhood NO_2_ exposure on global intracellular directional diffusion at the specified levels of sleep duration (i.e., slopes in Figure 1a were not significantly different from zero at 6, 7, and 8 hours of sleep). However, the significance of the interaction suggests a pattern of association between sleep duration and global intracellular directional diffusion may exist but at different durations of sleep (i.e., less than 6 hours or more than 8 hours). We additionally found that the positive relationship between prenatal O_3_ exposure and global white matter intracellular directional diffusion remained significant in those with lower sleep efficiency (i.e., 85%, 87%) but diminished as sleep efficiency increased. This suggests that higher sleep efficiency may buffer the brain’s white matter against the effects of prenatal O_3_ exposure.

Using RSI, intracellular directional diffusion in white matter likely represents diffusion within an axon – higher values may represent increased axon quantity, caliber, density, or myelination (44,46). Previous research has suggested air pollution in the prenatal period as well as later in childhood may influence white matter brain connectivity (17–21,23). Expanding upon these findings, in the current study, we found that those with longer sleep duration and higher sleep efficiency had lower global intra-axonal diffusion when exposed to certain noxious gaseous pollutants in the prenatal and childhood developmental periods. Regional analyses revealed that distinct commissural, association, and projection tracts (i.e., corpus callosum, uncinate fasciculus, and corticospinal tract) showed the strongest associations. Both the corticospinal tract and corpus callosum are vital for sensorimotor function (54,55). The uncinate fasciculus connects the amygdala and other parts of the temporal lobe to the medial orbitofrontal cortex, and while its functions are not entirely clear, it may be involved in emotional processing (56,57), behavioral inhibition (58), and impaired object naming (59). Alterations to the developmental trajectories of these tracts, either by attenuating or accelerating maturation, may impair learning and subsequent cognitive and emotional development (60,61).

Childhood NO_2_ exposure may cause neurotoxicity via the acute or chronic systemic inflammation it induces, beginning at the level of the lung alveoli (1,26). Upon inhalation, an innate immune reaction occurs in the lungs, whereby immune cells signal an upregulation of pro-inflammatory cytokines and induce oxidative stress, with immune components then passing into systemic circulation (1,26). This inflammatory cascade can contribute to BBB breakdown, leading to neuroinflammation and metal dyshomeostasis (10). Additionally, NO_2_ has been shown to contribute to mitochondrial dysfunction, which may be important in the context of white matter changes as it has been linked to oligodendrocyte damage (62,63). While the childhood pollutant exposure window (ages 9-10 years) is not completely concurrent with the available sleep and imaging data (ages 10-13 years) used in this study, there is evidence to suggest that annual averages are relatively stable prior to the year 2016, with more recent evidence from the U.S. EPA suggesting that concentrations remain relatively stable during the study period (2016–2020) (37–39,64). Our results indicate a significant interaction between childhood NO_2_ exposure and sleep duration, but it is not clear if this is beneficial to our brain outcome of interest given that the trends for the relationship between the pollutant and white matter microstructure were insignificant at the levels of sleep duration tested, as well as due to the cross-sectional nature of this analysis. This is consistent with previous work from our group demonstrating that childhood NO_2_ exposure was not related to intracellular directional diffusion in white matter cross-sectionally at ages 9-10 years nor longitudinally over a two-year follow-up period (19). However, we did find that childhood NO_2_ was negatively correlated with white blood cell counts, and that white blood cells counts were associated with changes in white matter microstructure in male youth at ages 10-13 years-old (2-year follow-up visit) in the ABCD Study (18). This may be indicative of possible acute or chronic changes/deficits in immune reactivity associated with childhood NO_2_ exposure. Longer sleep duration may aid in immune support and mitigate some of the negative effects of NO_2_ exposure, or it could indicate the presence of depressive symptomatology which may compound the pollutant’s toxic effects.

Here, we also find prenatal O_3_ exposure is related to higher white matter RND. Though exposure is from a different developmental window, this is consistent with previous work from our group using the ABCD Study dataset demonstrating that while there was a negative correlation between childhood O_3_ exposure and RND at age 9 in both sexes, higher childhood O_3_ exposure was associated with an accelerated increase in RND over time compared to those with less than average exposure (19). Given the prenatal exposure window in this current study, a plausible neurotoxic mechanism may be maternal oxidative stress and inflammation (both systemic and placental) (65). Inflammation and immune activation during pregnancy as a result of air pollution exposure has been linked to the onset of some neurodevelopmental disorders (i.e., autism spectrum disorder) (1,66), which have also been associated with hypermyelination in childhood (67,68). While the youth in this sample are unlikely have these neurodevelopmental phenotypes due to exclusion criteria at enrollment, it is possible that prenatal exposure to O_3_ contributes to hypermyelination at a subclinical level. A potential mechanism by which sleep efficiency may improve brain outcomes in the context of higher prenatal exposure to O_3_ includes through activity of neurotrophins like nerve growth factor (NGF) and brain-derived neurotrophic factor (BDNF). Prenatal exposure to O_3_ has been linked to decreased NGF in the hippocampus and increased BDNF in the striatum in a rodent model (69). As NGF has been shown to inhibit myelination in the CNS by oligodendrocytes (70) and BDNF has been shown to enhance myelination (71), prenatal exposure to O_3_ specifically may lead to hypermyelination in youth. The relationships between these neurotrophic factors and sleep are complex, but poor sleep has been linked to lower serum NGF in adolescents (72); thus, better sleep efficiency may increase NGF levels, potentially buffering against the effects of prenatal O_3_ on NGF and the resultant hypermyelination. In other words, higher sleep efficiency may result in higher NGF levels, thus aiding in the inhibition of aberrant CNS myelination in response to prenatal O_3_ exposure. However, additional work with multiple time points and markers of neurotrophic levels in the brain will be necessary to confirm these speculations.

There are several strengths and limitations associated in the current study. The question at hand, whether sleep (duration and efficiency) can modify the effects of ambient air pollution on structural brain connectivity, is novel and ultimately may help determine if sleep interventions could partially mitigate air pollution’s neurological effects in youth. Instead of using self-report questionnaire data, we used objective wearable-based measures of sleep duration and efficiency, reducing self-report bias (77). However, there are limitations to objective sleep measures from wearables like Fitbit Charge 2, such as subject compliance with protocol and inaccurate estimation of sleep duration and efficiency by Fitbit devices compared to polysomnography (78). Additionally, while we have pollutant concentration estimates at two different windows of developmental vulnerability, allowing us to characterize some differences in timing of exposure, there is currently no data available for pollutant concentrations concurrent with both the neuroimaging and sleep data when the children are ages 10-13 years. Future releases of ABCD Study datasets will eventually resolve this, and the results would be strengthened by examining air pollutant concentrations at this time point in addition to the two already included here. Additional limitations are those inherent to neuroimaging data, namely motion artifacts, which we accounted for by using only data that passed stringent quality control, had no clinically significant incidental findings, and by controlling for head motion within our models. Perhaps the biggest limitation to the current study is its cross-sectional nature - we only capture a snapshot of how sleep interacts with pollutant neurotoxicity, and future longitudinal studies will be able to more fully characterize how sleep affects brain developmental trajectories as they pertain to pollutant exposures. Additionally, while we show sleep metrics as moderating factors, poor sleep outcomes have also been associated with air pollution exposure (73) and may mediate the relationship between pollutants and brain outcomes. For instance, air pollution could feasibly impair brain waste clearance by inducing reactive astrogliosis, resulting in the swelling of astrocytic endfeet and impaired waste clearance through the perivascular spaces (74–76). Future studies are needed to disentangle these relationships, and longitudinal data will be especially important in determining how sleep may improve long-term resilience to neurotoxic pollutants. Lastly, the sample used here is large and regionally diverse, but not representative of the U.S. population or the larger ABCD Study cohort (79,80). Generally, the ABCD Study has an over-representation of subjects from wealthier and more educated backgrounds and an under-representation of Black and Asian participants. Additionally, Mroczek and colleagues (81) have voiced concerns regarding the overuse of publicly available datasets, in that multiple studies published using the same dataset may inflate the literature and contribute to issues of generalizability by perpetuating bias associated with sample nuances. Given this, these findings require validation in other diverse study populations. While the analysis provides valuable insights into the relationship between prenatal and childhood pollution exposure and brain outcomes, it is important to note that the study remains correlational in nature. Although controlling for demographic factors strengthens the findings by reducing potential confounding, the observational design of the study limits our ability to make definitive causal claims. To draw stronger causal inferences, further research employing more rigorous methods, such as randomized controlled trials or advanced causal inference techniques, will be necessary.

In conclusion, the current study demonstrates evidence that objective measures of sleep (i.e., duration and efficiency) interact with pollutant concentrations at two important windows of development to influence white matter microstructural integrity, despite the relatively low levels of pollutant exposure. Given sleep’s potential role in protecting young brains from neurotoxic air pollution in the face of a changing climate, encouraging healthy sleeping behaviors may help mitigate some of the negative neurotoxic effects of air pollution exposure in youth, thereby potentially increasing resilience to downstream behavioral outcomes.

## CRediT authorship contribution statement

**Devyn L. Cotter:** Writing – original draft, Writing – review & editing, Visualization, Formal analysis, Conceptualization. **Orsolya Kiss**: Writing – review & editing, Methodology. **Hedyeh Ahmadi:** Writing – review & editing, Methodology, Formal analysis. **Alethea de Jesus:** Writing – review & editing, Formal analysis. **Joel Schwartz:** Writing – review & editing, Methodology, Funding acquisition, Data curation. **Fiona C. Baker**: Writing – review & editing, Methodology, Data curation. **Daniel E. Hackman:** Methodology, Funding acquisition. **Megan M. Herting:** Writing – review & editing, Supervision, Resources, Project administration, Methodology, Funding acquisition.

## Acknowledgements

We would like to gratefully acknowledge Jade Li for her help with creating the brain tract visualizations.

