## Supplemental Material for "Sleep duration and efficiency moderate the effects of prenatal and childhood ambient pollutant exposure on global white matter microstructural integrity in adolescence"

**Supplemental Table 1.** Comparison of our sample to the entire ABCD cohort

| Cohort Characteristics | | |
|---|---|---|
| | Full Dataset (N=11876) | Subset for Analyses (N=2178) |
| Mean Age [months], (SD) | 144 (7.95) | 143.12 (7.72) |
| Sex [F], N (%) | 5680 (47.8%) | 995 (45.7%) |
| Race/Ethnicity, N (%) | | |
| Non-Hispanic White | 6182 (52.1%) | 1424 (65.4%) |
| Non-Hispanic Black | 1784 (15.0%) | 144 (6.6%) |
| Hispanic | 2411 (20.3%) | 367 (16.8%) |
| Non-Hispanic Asian | 252 (2.1%) | 38 (1.7%) |
| Multi-Racial/Other* | 1247 (10.5%) | 205 (9.4%) |
| Highest Household Education, N (%) | | |
| Post Graduate Degree | 2996 (25.2%) | 887 (40.7%) |
| Bachelor | 3338 (28.1%) | 660 (30.3%) |
| Some College | 3494 (29.4%) | 480 (22.1%) |
| HS Diploma/GED | 1261 (10.6%) | 111 (5.1%) |
| < HS Diploma | 787 (6.6%) | 40 (1.8%) |
| Overall Income (USD), N (%) | | |
| ≥100K | 4564 (38.4%) | 1057 (48.5%) |
| ≥50K & <100K | 3071 (25.9%) | 624 (28.7%) |
| <50K | 3223 (27.1%) | 384 (17.6%) |
| Don't Know/Refuse | 1016 (8.6%) | 113 (5.2%) |
| Missing | 2 | 0 |

**Supplemental Figure 1.** Spearman correlation plot between sleep and air pollution variables. $PM_{2.5}$ concentrations are in ug/m$^3$, while $NO_2$ and $O_3$ concentrations are in parts per billion (ppb). Sleep duration is in hours (h) and sleep efficiency is in percentages (%).

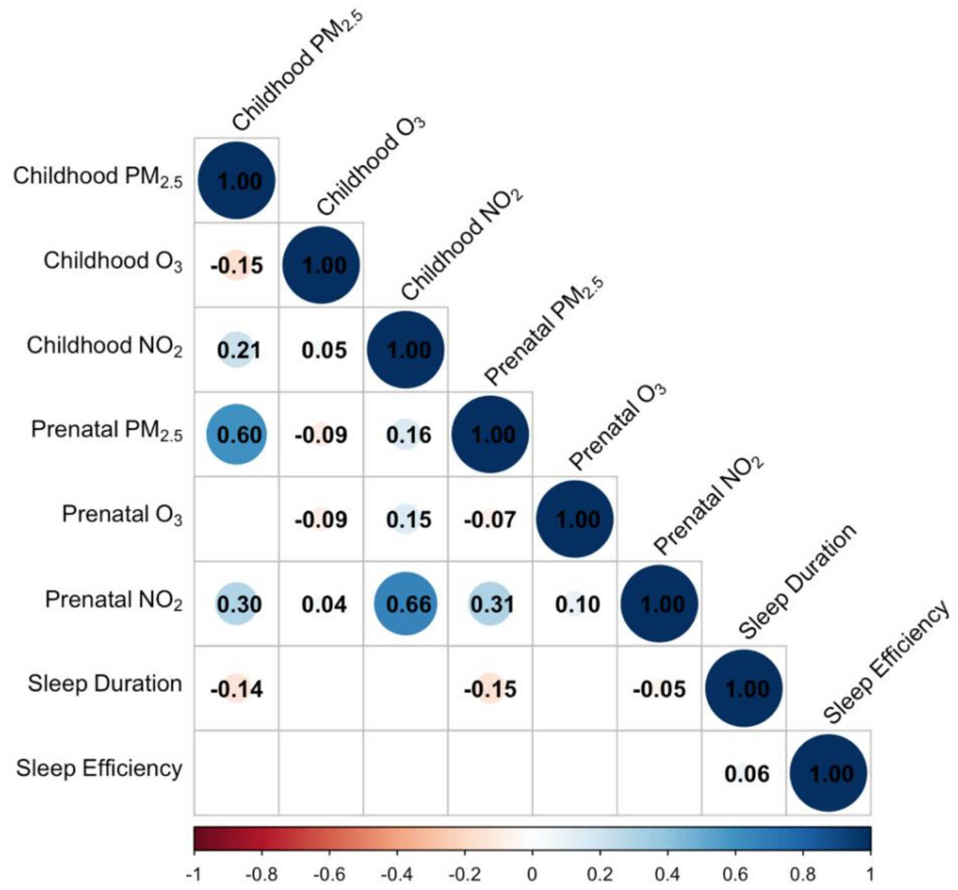

**Supplemental Figure 2.** Flow chart of participant exclusion.

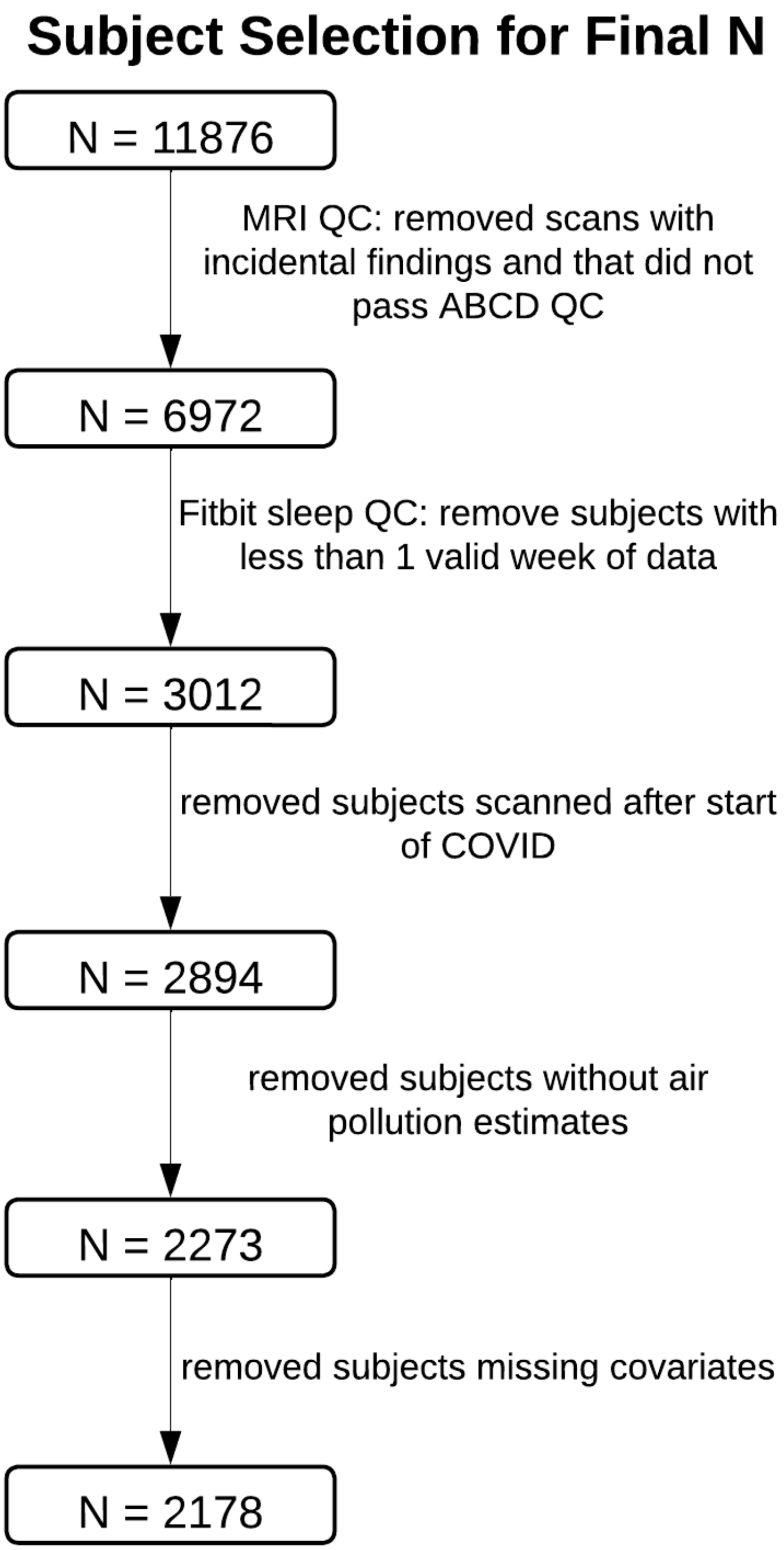

**Supplemental Table 2.** Estimated marginal means of linear trends for the childhood $NO_2$-by-sleep duration interaction term. Abbreviations: hours (h), standardized (std), standard error (SE), confidence interval (CI).

| Estimated marginal means of linear trends | | | | |
|---|---|---|---|---|
| Sleep Duration (h) | Childhood $NO_2$ (std) Slope | SE | Lower CI | Upper CI |
| 6 | 0.0013 | 0.0008 | -0.0002 | 0.0029 |
| 7 | 0.0005 | 0.0005 | -0.0005 | 0.0014 |
| 8 | -0.0004 | 0.0005 | -0.0014 | 0.0006 |
| Sleep Duration (h) Contrasts | Estimate | SE | z.ratio | p.value |
| 6-7 | **0.0009** | **0.0004** | **2.1320** | **0.0330** |
| 6-8 | **0.0018** | **0.0008** | **2.1320** | **0.0330** |
| 7-8 | **0.0009** | **0.0004** | **2.1320** | **0.0330** |

**Supplemental Table 3.** Estimated marginal means of linear trends for the prenatal $O_3$-by-sleep efficiency interaction term. Abbreviations: percentage (%), standardized (std), standard error (SE), confidence interval (CI).

| Estimated marginal means of linear trends | | | | |
|---|---|---|---|---|
| Sleep Efficiency (%) | Prenatal $O_3$ (std) Slope | SE | Lower CI | Upper CI |
| 0.86 | **0.0013** | **0.0005** | **0.0004** | **0.0022** |
| 0.87 | **0.0009** | **0.0004** | **0.0000** | **0.0018** |
| 0.88 | 0.0005 | 0.0005 | -0.0004 | 0.0014 |
| Sleep Efficiency (%) Contrasts | Estimate | SE | z.ratio | p.value |
| 0.86 – 0.87 | 0.0004 | 0.0002 | 2.4920 | 0.0127 |
| 0.86- 0.88 | 0.0008 | 0.0003 | 2.4920 | 0.0127 |
| 0.87 – 0.88 | 0.0004 | 0.0002 | 2.4920 | 0.0127 |

**Supplemental Table 4.** Results from post-hoc multi-pollutant models examining how sleep duration interacts with childhood NO2 to affect RND in individual white matter tracts, including unstandardized betas, standard error (SE), 95% confidence intervals (CI), and p-values. Significant FDR-corrected models are **bolded** ($p < 0.05$). Models were adjusted for pollutants not included in the interaction term, demographic and socioeconomic variables for each child, and precision MRI variables (see Methods). Abbreviations: intracellular directional diffusion (RND), frontal superior corticostriate (fSCS) and parietal superior corticostriate (pSCS), striatal to inferior frontal cortex (SIFC), anterior thalamic radiations (ATR), corticospinal tract (CST), temporal superior longitudinal fasciculi (tSLF), parietal superior longitudinal fasciculi (pSLF), inferior longitudinal fasciculi (ILF), inferior fronto-occipital fasciculi (IFO), inferior frontal to the superior frontal cortex (IFSFC), cingulate cingulum (CGC), parahippocampal cingulum (CGH), fornix (Fx), uncinate fasciculi (Unc), corpus callosum (CC), forceps minor (F.min), forceps major (F.maj), left (L), right (R).

| Tract | | Hem | Childhood $NO_2$ (std) | | | | Sleep Duration (h) | | | | Childhood $NO_2$ (std) x Sleep Duration (h) | | | | |
|---|---|---|---|---|---|---|---|---|---|---|---|---|---|---|---|
| | | | *Coefficient* | *SE* | *95% CI* | *p* | *Coefficient* | *SE* | *95% CI* | *p* | *Coefficient* | *SE* | *95% CI* | *p* | *FDR p* |
| Projection | fSCS | L | 0.0003 | 0.0038 | -0.0072, 0.0078 | 0.9373 | -0.0002 | 0.0009 | -0.0019, 0.0015 | 0.8239 | 0 | 0.0005 | -0.001, 0.001 | 0.954 | 0.9540 |
| | | R | 0.0038 | 0.0046 | -0.0053, 0.0129 | 0.4093 | -0.0001 | 0.0009 | -0.0018, 0.0016 | 0.9071 | -0.0005 | 0.0006 | -0.0017, 0.0007 | 0.4182 | 0.5848 |
| | pSCS | L | 0.0029 | 0.0032 | -0.0034, 0.0091 | 0.3704 | 0.001 | 0.0008 | -0.0006, 0.0026 | 0.2158 | -0.0003 | 0.0004 | -0.0011, 0.0005 | 0.4716 | 0.5848 |
| | | R | 0.0038 | 0.004 | -0.0041, 0.0117 | 0.3478 | 0.0007 | 0.0009 | -0.001, 0.0024 | 0.4303 | -0.0005 | 0.0005 | -0.0015, 0.0005 | 0.3327 | 0.5157 |
| | SIFC | L | 0.0058 | 0.0047 | -0.0035, 0.0151 | 0.22 | -0.0009 | 0.0009 | -0.0026, 0.0008 | 0.2872 | -0.0008 | 0.0006 | -0.002, 0.0005 | 0.2257 | 0.4373 |
| | | R | 0.0047 | 0.0048 | -0.0048, 0.0141 | 0.3336 | -0.0006 | 0.0008 | -0.0022, 0.001 | 0.4441 | -0.0006 | 0.0006 | -0.0019, 0.0006 | 0.3125 | 0.5157 |
| | ATR | L | 0.0057 | 0.0049 | -0.0039, 0.0153 | 0.247 | -0.0004 | 0.0009 | -0.0021, 0.0013 | 0.6106 | -0.0009 | 0.0006 | -0.0021, 0.0003 | 0.1532 | 0.3667 |
| | | R | 0.0018 | 0.0045 | -0.0069, 0.0106 | 0.6813 | -0.0011 | 0.001 | -0.003, 0.0009 | 0.2735 | -0.0003 | 0.0006 | -0.0014, 0.0008 | 0.6095 | 0.6998 |
| | CST | L | -0.0003 | 0.0006 | -0.0014, 0.0008 | 0.6095 | 0.0002 | 0.0006 | -0.001, 0.0014 | 0.6946 | -0.0004 | 0.0005 | -0.0013, 0.0005 | 0.3308 | 0.5157 |
| | | R | 0.0008 | 0.0038 | -0.0066, 0.0082 | 0.8412 | 0.0001 | 0.0006 | -0.001, 0.0011 | 0.9239 | -0.0001 | 0.0005 | -0.0011, 0.0008 | 0.7603 | 0.7856 |
| Association | tSLF | L | 0.0045 | 0.0033 | -0.0019, 0.0108 | 0.169 | -0.0002 | 0.0006 | -0.0013, 0.0009 | 0.7609 | -0.0006 | 0.0004 | -0.0013, 0.0002 | 0.1608 | 0.3667 |
| | | R | 0.0036 | 0.0047 | -0.0055, 0.0128 | 0.4366 | -0.0002 | 0.0006 | -0.0014, 0.001 | 0.7227 | -0.0004 | 0.0006 | -0.0016, 0.0008 | 0.4992 | 0.5952 |
| | pSLF | L | 0.0036 | 0.0036 | -0.0034, 0.0107 | 0.3123 | -0.0001 | 0.0006 | -0.0013, 0.001 | 0.8374 | -0.0005 | 0.0005 | -0.0014, 0.0004 | 0.3183 | 0.5157 |
| | | R | 0.0069 | 0.0037 | -0.0003, 0.0141 | 0.0611 | 0 | 0.0005 | -0.0011, 0.0011 | 0.9947 | -0.0009 | 0.0005 | -0.0018, 0.0001 | 0.0714 | 0.2767 |
| | ILF | L | 0.0061 | 0.0042 | -0.0023, 0.0144 | 0.1532 | 0.0005 | 0.0007 | -0.0009, 0.0018 | 0.525 | -0.0007 | 0.0005 | -0.0017, 0.0003 | 0.1656 | 0.3667 |
| | | R | 0.0069 | 0.0051 | -0.003, 0.0168 | 0.1734 | 0.0008 | 0.0007 | -0.0006, 0.0021 | 0.2714 | -0.0008 | 0.0006 | -0.0021, 0.0004 | 0.2024 | 0.4183 |
| | IFO | L | *0.0077* | *0.0035* | 0.0008, 0.0146 | *0.0296* | -0.0004 | 0.0006 | -0.0016, 0.0008 | 0.5223 | *-0.0011* | *0.0005* | -0.002, -0.0001 | *0.0238* | *0.1476* |
| | | R | 0.0037 | 0.0026 | -0.0014, 0.0089 | 0.1564 | -0.0005 | 0.0005 | -0.0015, 0.0004 | 0.2794 | -0.0005 | 0.0003 | -0.0012, 0.0001 | 0.1123 | 0.3517 |
| | IFSFC | L | -0.0013 | 0.0038 | -0.0088, 0.0062 | 0.7421 | 0.0004 | 0.0009 | -0.0013, 0.0021 | 0.6466 | 0.0002 | 0.0005 | -0.0008, 0.0012 | 0.6651 | 0.7364 |
| | | R | 0.0026 | 0.0036 | -0.0045, 0.0096 | 0.4733 | 0.0003 | 0.0007 | -0.0011, 0.0017 | 0.7005 | -0.0004 | 0.0004 | -0.0012, 0.0005 | 0.4224 | 0.5848 |
| Assoc - Limbic | CGC | L | 0.0101 | 0.0064 | -0.0024, 0.0226 | 0.1138 | -0.0005 | 0.0009 | -0.0022, 0.0013 | 0.589 | -0.0013 | 0.0008 | -0.0029, 0.0004 | 0.1248 | 0.3517 |
| | | R | 0.0034 | 0.0088 | -0.014, 0.0207 | 0.7021 | -0.0009 | 0.0017 | -0.0042, 0.0024 | 0.5916 | -0.0004 | 0.0012 | -0.0027, 0.0019 | 0.7528 | 0.7856 |
| | CGH | L | 0.0094 | 0.0056 | -0.0015, 0.0204 | 0.0917 | *0.0028* | *0.0012* | 0.0005, 0.0052 | *0.0171* | -0.0011 | 0.0007 | -0.0024, 0.0003 | 0.1175 | 0.3517 |
| | | R | *0.013* | *0.0052* | 0.0029, 0.0232 | *0.0119* | *0.003* | *0.0009* | 0.0012, 0.0048 | *0.001* | *-0.0015* | *0.0007* | -0.0029, -0.0001 | *0.0343* | *0.1772* |
| | Fx | L | 0.0038 | 0.0043 | -0.0048, 0.0123 | 0.3866 | -0.0008 | 0.0013 | -0.0033, 0.0017 | 0.5363 | -0.0004 | 0.0006 | -0.0015, 0.0007 | 0.4695 | 0.5848 |
| | | R | 0.0031 | 0.0047 | -0.0061, 0.0124 | 0.5046 | -0.0007 | 0.0011 | -0.0029, 0.0015 | 0.5248 | -0.0005 | 0.0007 | -0.0018, 0.0008 | 0.4652 | 0.5848 |
| | Unc | L | 0.0117 | 0.0062 | -0.0004, 0.0239 | 0.0582 | -0.0006 | 0.0009 | -0.0025, 0.0012 | 0.4992 | -0.0016 | 0.0008 | -0.0032, 0.0001 | 0.0621 | 0.2750 |
| | | R | *0.0133* | *0.0032* | 0.007, 0.0195 | *0* | 0.0004 | 0.0006 | -0.0008, 0.0015 | 0.5497 | ***-0.0018*** | ***0.0004*** | -0.0027, -0.001 | ***0.00002*** | ***0.0006*** |
| Commissural | CC | BL | *0.0086* | *0.0025* | 0.0037, 0.0135 | *0.0006* | -0.0004 | 0.0007 | -0.0018, 0.0009 | 0.5379 | ***-0.0012*** | ***0.0003*** | -0.0018, -0.0006 | ***0.0002*** | ***0.0031*** |
| | f.min | BL | *0.0091* | *0.004* | 0.0014, 0.0169 | *0.0209* | -0.0009 | 0.001 | -0.003, 0.0011 | 0.3595 | *-0.0012* | *0.0005* | -0.0022, -0.0002 | *0.0168* | *0.1476* |
| | f.maj | BL | *0.0109* | *0.0047* | 0.0017, 0.0201 | *0.0203* | -0.0002 | 0.0011 | -0.0023, 0.0019 | 0.8298 | *-0.0015* | *0.0006* | -0.0027, -0.0002 | *0.0227* | *0.1476* |

**Supplemental Table 5.** Results from post-hoc multi-pollutant models examining how sleep duration interacts with prenatal O3 to affect RND in individual white matter tracts, including unstandardized betas, standard error (SE), 95% confidence intervals (CI), and p-values. Significant FDR-corrected models are **bolded** ($p < 0.05$). Models were adjusted for pollutants not included in the interaction term, demographic and socioeconomic variables for each child, and precision MRI variables (see Methods). Abbreviations: intracellular directional diffusion (RND), frontal superior corticostriate (fSCS) and parietal superior corticostriate (pSCS), striatal to inferior frontal cortex (SIFC), anterior thalamic radiations (ATR), corticospinal tract (CST), temporal superior longitudinal fasciculi (tSLF), parietal superior longitudinal fasciculi (pSLF), inferior longitudinal fasciculi (ILF), inferior fronto-occipital fasciculi (IFO), inferior frontal to the superior frontal cortex (IFSFC), cingulate cingulum (CGC), parahippocampal cingulum (CGH), fornix (Fx), uncinate fasciculi (Unc), corpus callosum (CC), forceps minor (F.min), forceps major (F.maj), left (L), right (R).

| Tract | | Hem | Prenatal $O_3$ (std) | | | | Sleep Efficiency (%) | | | | Prenatal $O_3$ (std) x Sleep Efficiency (%) | | | | |
|---|---|---|---|---|---|---|---|---|---|---|---|---|---|---|---|
| | | | *Coefficient* | *SE* | *95% CI* | *p* | *Coefficient* | *SE* | *95% CI* | *p* | *Coefficient* | *SE* | *95% CI* | *p* | *FDR p* |
| Projection | fSCS | L | 0.0152 | 0.0151 | -0.0145, 0.0449 | 0.316 | 0.0128 | 0.0194 | -0.0254, 0.0509 | 0.5117 | -0.0168 | 0.0176 | -0.0513, 0.0176 | 0.3382 | 0.5459 |
| | | R | 0.0131 | 0.0206 | -0.0273, 0.0534 | 0.5264 | -0.0107 | 0.0209 | -0.0517, 0.0302 | 0.6066 | -0.0139 | 0.0239 | -0.0608, 0.0331 | 0.5623 | 0.7263 |
| | pSCS | L | 0.0193 | 0.0201 | -0.0202, 0.0588 | 0.3383 | *0.0359* | *0.0175* | 0.0015, 0.0703 | *0.0409* | -0.0209 | 0.0232 | -0.0664, 0.0246 | 0.367 | 0.5459 |
| | | R | 0.0291 | 0.0209 | -0.0119, 0.07 | 0.164 | -0.0109 | 0.0204 | -0.051, 0.0292 | 0.5949 | -0.0313 | 0.024 | -0.0785, 0.0158 | 0.1921 | 0.3722 |
| | SIFC | L | 0.0348 | 0.0251 | -0.0145, 0.0841 | 0.1662 | -0.021 | 0.0226 | -0.0654, 0.0234 | 0.3532 | -0.0392 | 0.0286 | -0.0954, 0.0169 | 0.1708 | 0.3530 |
| | | R | 0.0058 | 0.0171 | -0.0278, 0.0395 | 0.7333 | -0.01 | 0.0243 | -0.0577, 0.0377 | 0.6809 | -0.0061 | 0.0197 | -0.0447, 0.0325 | 0.756 | 0.8370 |
| | ATR | L | 0.0369 | 0.0197 | -0.0017, 0.0755 | 0.0611 | -0.0154 | 0.0189 | -0.0526, 0.0217 | 0.4156 | -0.0425 | 0.0226 | -0.0867, 0.0018 | 0.0598 | 0.1823 |
| | | R | 0.0033 | 0.0175 | -0.031, 0.0377 | 0.8487 | 0.0188 | 0.0207 | -0.0217, 0.0593 | 0.3634 | -0.0033 | 0.0203 | -0.0431, 0.0364 | 0.869 | 0.8998 |
| | CST | L | *0.0344* | *0.0162* | 0.0026, 0.0662 | *0.0342* | 0.0125 | 0.0153 | -0.0176, 0.0425 | 0.4167 | *-0.0394* | *0.0187* | -0.0761, -0.0028 | *0.0348* | *0.1823* |
| | | R | *0.0366* | *0.0096* | 0.0177, 0.0555 | *0.0001* | -0.0146 | 0.0086 | -0.0315, 0.0023 | 0.0914 | ***-0.0412*** | ***0.0114*** | -0.0635, -0.0189 | ***0.0003*** | ***0.0093*** |
| Association | tSLF | L | *0.0425* | *0.0212* | 0.0009, 0.0842 | *0.0451* | 0.0028 | 0.0192 | -0.0349, 0.0405 | 0.8848 | *-0.048* | *0.0243* | -0.0957, -0.0003 | *0.0484* | *0.1823* |
| | | R | *0.0288* | *0.0138* | 0.0018, 0.0559 | *0.0368* | -0.0095 | 0.0125 | -0.034, 0.0149 | 0.4442 | *-0.0325* | *0.0157* | -0.0634, -0.0017 | *0.0389* | *0.1823* |
| | pSLF | L | 0.046 | 0.0246 | -0.0022, 0.0942 | 0.0612 | 0.0129 | 0.0229 | -0.0321, 0.0579 | 0.5744 | -0.0519 | 0.0281 | -0.1069, 0.0032 | 0.0647 | 0.1823 |
| | | R | *0.0375* | *0.0188* | 0.0006, 0.0745 | *0.0467* | -0.0089 | 0.0184 | -0.0449, 0.0271 | 0.6274 | *-0.0425* | *0.0215* | -0.0846, -0.0004 | *0.0479* | *0.1823* |
| | ILF | L | 0.0174 | 0.0193 | -0.0203, 0.0552 | 0.3655 | 0.0176 | 0.02 | -0.0215, 0.0568 | 0.377 | -0.0192 | 0.0223 | -0.0629, 0.0245 | 0.3894 | 0.5459 |
| | | R | 0.0197 | 0.0184 | -0.0165, 0.0558 | 0.2857 | 0.0079 | 0.0209 | -0.0332, 0.0489 | 0.7075 | -0.0212 | 0.0211 | -0.0627, 0.0202 | 0.3143 | 0.5459 |
| | IFO | L | *0.047* | *0.018* | 0.0117, 0.0823 | *0.009* | -0.0111 | 0.021 | -0.0523, 0.0301 | 0.5964 | *-0.0532* | *0.0208* | -0.094, -0.0123 | *0.0108* | *0.1147* |
| | | R | *0.0287* | *0.0138* | 0.0017, 0.0558 | *0.0375* | 0.0054 | 0.0171 | -0.0281, 0.039 | 0.7508 | *-0.0316* | *0.0158* | -0.0626, -0.0007 | *0.0452* | *0.1823* |
| | IFSFC | L | 0.0107 | 0.0217 | -0.0318, 0.0532 | 0.6224 | 0.0147 | 0.0223 | -0.029, 0.0583 | 0.5094 | -0.0108 | 0.0248 | -0.0595, 0.0378 | 0.6626 | 0.7900 |
| | | R | 0.036 | 0.0207 | -0.0046, 0.0767 | 0.0825 | -0.0231 | 0.0165 | -0.0555, 0.0093 | 0.1624 | -0.0403 | 0.024 | -0.0875, 0.0068 | 0.0936 | 0.2073 |
| Assoc - Limbic | CGC | L | 0.0337 | 0.0385 | -0.0419, 0.1092 | 0.3823 | -0.0391 | 0.036 | -0.1097, 0.0315 | 0.2775 | -0.0378 | 0.044 | -0.124, 0.0485 | 0.3904 | 0.5459 |
| | | R | 0.0047 | 0.0354 | -0.0647, 0.0742 | 0.8939 | 0.0379 | 0.0396 | -0.0397, 0.1155 | 0.338 | -0.0054 | 0.0404 | -0.0847, 0.0739 | 0.8935 | 0.8998 |
| | CGH | L | 0.0362 | 0.0398 | -0.0419, 0.1142 | 0.3637 | 0.0567 | 0.035 | -0.0119, 0.1252 | 0.1051 | -0.0398 | 0.0456 | -0.1292, 0.0497 | 0.3833 | 0.5459 |
| | | R | 0.03 | 0.036 | -0.0406, 0.1007 | 0.4048 | 0.0706 | 0.0404 | -0.0086, 0.1499 | 0.0807 | -0.0348 | 0.0417 | -0.1166, 0.0471 | 0.405 | 0.5459 |
| | Fx | L | -0.0082 | 0.0196 | -0.0467, 0.0302 | 0.6744 | 0.0047 | 0.0259 | -0.046, 0.0554 | 0.8553 | 0.0111 | 0.0225 | -0.0329, 0.0552 | 0.6201 | 0.7689 |
| | | R | -0.0073 | 0.0222 | -0.0509, 0.0363 | 0.7425 | -0.036 | 0.0294 | -0.0936, 0.0216 | 0.2205 | 0.0086 | 0.0256 | -0.0417, 0.0589 | 0.7369 | 0.8370 |
| | Unc | L | 0.0529 | 0.0289 | -0.0039, 0.1096 | 0.0678 | -0.0065 | 0.0312 | -0.0677, 0.0547 | 0.8345 | -0.0596 | 0.0331 | -0.1245, 0.0054 | 0.0723 | 0.1868 |
| | | R | 0.0291 | 0.0148 | 0.0001, 0.0582 | 0.0495 | 0.0071 | 0.0251 | -0.0422, 0.0564 | 0.7768 | -0.0323 | 0.0173 | -0.0663, 0.0016 | 0.0621 | 0.1823 |
| Commissural | CC | BL | 0.0287 | 0.016 | -0.0026, 0.06 | 0.072 | -0.0031 | 0.0156 | -0.0337, 0.0276 | 0.8452 | -0.0318 | 0.0184 | -0.068, 0.0043 | 0.0839 | 0.2001 |
| | f.min | BL | *0.0577* | *0.0222* | 0.0142, 0.1012 | *0.0094* | -0.0165 | 0.0195 | -0.0548, 0.0218 | 0.3978 | *-0.0652* | *0.0256* | -0.1154, -0.0149 | *0.0111* | *0.1147* |
| | f.maj | BL | 0.0051 | 0.0259 | -0.0457, 0.0559 | 0.8449 | 0.0077 | 0.0238 | -0.0389, 0.0544 | 0.7447 | -0.0037 | 0.0296 | -0.0617, 0.0543 | 0.8998 | 0.8998 |
